# Genetic diversity and haplotypic structure of a Saudi population sample using Investigator Argus X-12 amplification kit

**DOI:** 10.1101/760819

**Authors:** Safia A. Messaoudi, Saranya R. Babu, Abrar B. Alsaleh, Mohamed Albajjah, Noora AlSnan, Abdul Rauf Chaudhary

**Author notes:** Corresponding Author: Dr Safia A. Messaoudi, Forensic Biology Department, Naif Arab University for Security Sciences, P.O Box: 6830 Riyadh 11452, Kingdom of Saudi Arabia. **Co-Authors emails:** Saranya Babu Abrar Alsaleh Mohamed Albajja Noora Al Snan Abdul Rauf Chaudhary.

## Abstract

X-chromosome short tandem repeat (X-STR) markers have shown a great capability in forensic identity investigations and paternity testing involving kinship analysis.

In the current study, the distribution of 12 X-STR loci located in four linkage groups was evaluated using Investigator^®^ Argus X-12 Amplification Kit in 200 unrelated healthy individuals (105 males and 95 females) from the central region of the Saudi Arabia in order to create a DNA database.

Our results indicated that DXS10146 locus was the most informative with 21 alleles while DXS8378 locus was the least with five alleles. Forensic parameters showed that all X-STRs loci either as individual markers or as linkage groups provide genetic information with high discrimination that is appropriate for forensic purposes with Paternity Informed Consent (PIC), Power of exclusion (PE), and Paternity index (PI) varied from 0.61211 to 0.917979, 0.38722 to 0.842949, and 0.038416 to 0.16367, respectively. A significant Linkage disequilibrium (LD) with *p*-value after Bonferroni correction *p* ≤ 0.05/66= 0.0008 was observed for 17 pairs of loci in male samples and 4 pairs of loci in female. In the male group, LG3 showed relatively high values of Haplotype diversity. The pairwise genetic distance fixation index (Fst) results showed that the Saudi population is genetically close to the Egyptian and Emirati populations and distant to the Turkish population.

The current study revealed that Investigator® Argus 12 X-STR kit would support forensic application, kinship testing involving female offspring, and human identification in Saudi populations.

## Introduction

The use of X-chromosome short tandem repeat (X-STR) markers has been greatly increased in the forensic setting over the last decade [1]. They have shown great forensic ability and are, therefore, recognized as efficient tools in forensic investigations of kinship cases [2–4]. A variety of X-STR systems have been developed and added to the list of conventional markers for better results in complex kinship analysis and cases involving mother-son and father/daughter relationship [5–7].

This forensic and human identity testing potential of X-STRs is due to the inheritance pattern of the X-chromosome compared to other genetic markers [2]. Any forensically questioned relationship, where at least one female is involved, may benefit from the use of X-STRs which can be applied to cases of missing persons, criminal incest, immigration, deficiency paternity or other similar situation [8].

Since the X-chromosome recombines only in females [5], therefore, X-STRs show stronger linkage disequilibrium (LD) than autosomal STRs.

Szibor and Hering [5] described four original linkage groups contained the following markers: DXS8378 and DXS9902 in linkage group 1; DXS7132, DXS6789, DXS101, DXS7424, and GATA172D05 in linkage group 2; HPRTB in linkage group 3; and DXS7423 in linkage group 4.

LD between X-STR loci has proven to be population-specific and is further dependent on factors such as random drift, founder effect, mutation rates, selection, and population admixture [9,10].

Among haploid markers and in addition to STRs, stable haplotypes of closely associated X-chromosome markers have proven to be a powerful, highly informative and particularly profitable tool in the analysis of complex kinship cases [11,2].

Saudi Arabia is the largest country in the Arabian Peninsula with a population exceeding 32 million where people are non-uniformly distributed [12]. Nearly half of the population resides in two major administrative provinces, Riyadh and Mecca [13]. Indigenous Arab people who seem to constitute ∼63% of the population are historically organized into geographically-differentiated patrilineal descent groups, or tribes [14], with a known tradition of consanguinity [15]. In addition, African and Asian surrounding populations have also influenced the genetic structure of Arab population [16,17].

The majority of Saudi Arabian Y-chromosome composition has been reported to be of Levantine origins (69%); with significant contributions from the East via Iran (17%), and Africa (14%) [17].

This geographical and social organization might be expected to have an effect on patterns of genetic diversity. However, allele/haplotype frequency data of forensically relevant X-STR markers in Saudi Arabia are extremely limited.

The present study is the first to characterize X-STRs-based genetic diversity of Saudi Arabian population from the Central region using Investigator™ Argus X-12 amplification kit. Twelve gonosomal STR markers were studied to estimate the allelic frequencies and evaluate different forensic and genetic population parameters in a sample of 105 unrelated 105 Saudi male and 95 female volunteers. It is hoped that present investigations will contribute significantly to the existing state of knowledge regarding the X-STR genetic diversity in Saudi population.

## Materials and methods

The present study was undertaken to determine allele frequencies and statistical parameters of forensic interest of 12 X-STR loci (DXS7132, DXS7423, DXS8378, DXS10074, DXS10079, DXS10101, DXS10103, DXS10134, DXS10135, DXS10146, DXS10148, and HPRTB) in a randomly selected cohort of unrelated Saudi individuals from the central region of the kingdom of Saudi Arabia for the purpose of constructing closely linked STR markers groups on the X-chromosome.

### Samples collection

Buccal swabs were obtained from 200 natives (up to three generations), unrelated healthy adult, fully informed and consented male (n=105) and female (n=95) volunteers living in the central region of Saudi Arabia. Demographic and past medical history was collected through a standard questionnaire. Individuals having documented genetic disorders, bone marrow transplant, and recent whole blood transfusion were excluded. The study was approved by the Ethical Committee for Scientific Research at the Naif Arab University of Security Sciences, Riyadh, Saudi Arabia.

### DNA extraction, amplification and fragment detection

Genomic DNA was isolated from buccal swabs using the Chelex extraction method [18]. Extracted DNA samples were quantified following the Quantifiler Duo Human DNA Quantification Kit (Thermo Fisher Scientific, Inc., Waltham, MA, USA) as recommended by the manufacturer in 7500 Real-time PCR System (Thermo Fisher Scientific Company, Carlsbad, USA) to normalize the input DNA quantity in multiplex PCR reaction.

The 12 X-STR loci were simultaneously amplified by the multiplex PCR using the Investigator Argus X-12 Amplification Kit (QIAGEN). Multiplex PCR was performed on an Applied Biosystems Veriti Thermal Cycler ((Thermo Fisher Scientific, Inc., Waltham, MA, USA)) in a 10.6 μl reaction volume containing 10 µL of Hi-Di formamide, with appropriate 0.1 µL size standard and 0.5 µL of the amplified PCR product. The samples were denatured for 3 min at 95°C and snap cooled on ice before loading onto the instrument. Amplified fragments were separated by capillary electrophoresis using 3130 Genetic Analyzer ((Thermo Fisher Scientific, Inc., Waltham, MA, USA), following the manufacturer’s instructions and size separated with internal lane size standard BTO-5 following manufacturer’s protocols (Qiagen N.V., Venlo, Netherlands).

Alleles were assigned according to the International Society of Forensic Genetics (ISFG) guidelines for forensic STR by comparing to the reference allelic ladder included in the kit using GeneMapper ID-X software v.1.2 (Thermo Fisher Scientific, Inc., Waltham, MA, USA). A peak detection threshold of 50 RFUs was used for allele designation.

### Forensic and population genetic parameters

Allele frequencies and haplotype frequencies were estimated for each locus using StatsX v2.0 software [19]. Fisher’s exact tests to evaluate the Hardy–Weinberg equilibrium (HWE) by locus and linkage disequilibrium (LD) between pair of loci were estimated with STRAF - a convenient online tool for STR data evaluation in forensic genetics [20]. Mean exclusion chance in Duos (MECD), PD for females (PDf), and PD in males (PDm) were estimated using ChrX-STR.org 2.0 website [21].

Interpopulation pairwise genetic distances based on Fst between central region Saudi population and the rest of populations extracted from the literature were calculated using POPTREE2 software [22] and represented by a nonmetric multidimensional scaling (NM-MDS) analysis using IBM SPSS Statistics v21.0 Software. Phylogenetic tree was constructed from allele frequency data by using the neighbour-joining method [23] *via* MEGA X: Molecular Evolutionary Genetics Analysis [24].

## Results and Discussion

### Intrapopulation Genetic diversity and Forensic efficiency

In the present study, allele frequencies, observed heterozygosity (Ho), expected heterozygosity (He) and *p*-value for Hardy-Weinberg equilibrium (HWE) were determined for the twelve X-chromosomal Short Tandem Repeat (STR) markers DXS8378, DXS10146, DXS7423, DXS10134, HPRTB, DXS10101, DXS10103, DXS10079, DXS10074, DXS7132, DXS10148 and DXS10135 within DNA samples from the population of central Saudi Arabia.

Out of 200 samples, 146 unique alleles were observed with the allele frequencies ranging from 0.0034 to 0.4542. Allele frequencies for each of the 12 X-STR markers in the population groups (male, female) are shown in Table 1. There were no significant differences in the distribution of allele frequencies between male and female samples (*p* > 0.05) using the exact test, which proved that these X-STR loci had no gender bias of allele distribution in the studied group. Therefore, we pooled male and female samples together for calculating forensic parameters.

The most polymorphic locus was DXS10146 with 21 alleles while DXS8378 was the least polymorphic locus with 5 alleles. The highest Gene Diversity (GD) was observed for locus DXS10135 with 0.9937 while the smallest for DXS7423 with a GD value of 0.661355 (Table 1)

HWE test was performed on female samples. All loci were in HWE, in the female Saudi population, with the exception of DXS10074, DXS10101, DXS10135 and DXS10148 even after Bonferroni correction (p > 0.05/12 = 0.0042) (Table 1). This deviation could be due to the consanguinity high rate and the smaller sample size.

Based on allele frequencies of the pooled data, we further determined the statistical parameters of forensic interest. The most informative locus was DXS10135 with 20 alleles while DXS8378 was lowest with 5 alleles. For the present studied population, the PIC, PE, and PI varied from 0.61211 to 0.917979, 0.38722 to 0.842949, and 0.038416 to 0.16367, respectively (Table 1).

The combined values of each forensic parameter, _c_PD_M_, _c_PD_F_, _c_MECK Rüger, cMEC Kishida, and cMEC Desmarais as well as _c_MEC Desmarais Duo, were 0.999999987962839, 0.999999999999998, 0.999999701236302, 0.999999972467329, 0.999999987434766 and 0.999999813797358, respectively (Table 1).

The present results indicate that all X-STRs loci either as individual markers or as linkage groups provide genetic information with high discrimination that is appropriate for forensic purposes. The high values of combined PDm and combined PDf, as well as high values of combined MEC in deficiency cases, normal trios, and duo indicated that these 12 X-STR loci of the Investigator® Argus X-12 kit would be suitable package as a complement of autosomal STRs for forensic identity and paternity testing involving kinship analysis when the offspring is female or investigating father-daughter relationships in the Saudi population.

### Linkage disequilibrium analysis and Haplotypic structure

Tests of linkage disequilibrium (LD) were performed for female and male samples for all pairs of loci in the group and the results were presented in Table 2 and 3. Significant LD with *p*-value ≤ 0.05 was observed for 19 pairs of loci in female and 29 pairs of loci in male samples. Moreover, and in order to improve the level of significance, Bonferroni correction was applied and significant results with *p* ≤0.05/66 (0.0008) were observed in 4 pairs of loci in female; DXS10103-DXS10148, DXS10146-DXS7132, HPRTB-DXS10079 and HPRTB-DXS10134 and 17 pairs of loci in male; DXS7132 showed the highest number of LD with the following 6 markers: DXS8378, DXS7423, DXS10103, DXS10079, DXS10135 and DXS10148, while DXS10134 showed LD with only one marker DXS10074. DXS7423 and HPRTB loci were in LD with DXS10146, DXS10148, DXS10079, DXS10103 and DXS10148, DXS10101, DXS8378, respectively. DXS8378 was in LD with DXS10148 and DXS10101 as well as with DXS7132 and HPRTB. The 12 X-STR markers are clustered into four linkage groups based on their physical localization [5,7]: LG1 (Xp22) contains DXS8378-DXS10135-DXS10148, LG2 (Xq11) contains DXS7132-DXS10074-DXS10079, LG3 (Xq26) contains DXS10101-DXS10103-HPRTB and LG4 (Xq28) contains DXS7423-DXS10134-DXS10146.

In male group, a significant association was identified between the pairwise LD and the markers located in the same LG (Table 3). The tight linkage between the X-STR trios in the four linkage groups are as follows: LG1 (DXS10148 and DXS8378), LG2 (DXS7132 and DXS10079), LG3 (DXS10101 and HPRTB) and LG4 (DXS7423 and DXS10146).

Whereas, no pairwise LD was found among the four LGs for the female group (Table 2). The lack of association between some of the markers inside linkage groups has advantages as it contributes to the high PDs value of this multiplex in the present population [1].

Linkage expectations were only based on physical distances between loci, but LD may also result from random genetic drift, founder effects, mutations, selection and population admixture or stratification [25,26].

Previous studies have demonstrated that loci inside each of the four X-STR linkage groups show reduced, but non-zero recombination rates [27]. In addition, recombination between linkage groups is less than 50% [7,28,29]. In fact, Diegoli and Rohde [7] argues against a simple model of independent X chromosomal linkage groups for these reasons.

Haplotype diversity [30] of four LGs composed of the closely linked X-STRs were tested in the male group (n=105 samples) and listed in Table 4. Each cluster of three X-STR markers is considered as one haplotype. The numbers of observed haplotypes in each of the 4 linkage groups-LG1, LG2, LG3, LG4 were 70, 69, 81 and 69, respectively, while HD values were 0.99063, 0.99042, 0.99353, and 0.990787, respectively.

The LG3 showed relatively high values of HD compared to other LGs. This indicated that these linkage groups were quite informative in the studied Saudi group and would have high application value in forensic sciences.

The three most common haplotype for LG1 was 12-29-26.1 displaying 4.76 % of haplotype frequency, in LG2 two sets of haplotypes 12-8-20 and 16-15-20 were observed each with a frequency of 3.8%. LG3 and LG4 presented haplotypes 29.2-19-13 and 15-34-45.2 with 3.8% and 4.76% of haplotype frequency respectively.

### Interpopulation diversity

To our knowledge, this is the first report from Saudi Arabia on these 12 markers, so no Saudi data is available for comparison. In the present study, Saudi allele frequencies were compared with eight worldwide populations from the literature to elucidate the genetic relationships between Saudis, Emiratis [31], Egyptians [32], Turkish [33], Algerians [34], Jewish [35], Filipino [36], Bangladeshis [37] and Indians [38] using the population pairwise genetic distance fixation index (Fst). Fst was used to perform the comparison, with differences being statistically significant for *p*-values < 0.001 after Bonferroni correction (Table 5).

The results of population Fst for 12 X-STRs was substantial, ranging from 0.004 to 0.052 indicating differences in allele frequencies between the studied populations that varied geographically, linguistically and socially.

The comparison with published data showed that Saudi population in this study had lower pairwise Fst values with those populations that are geographically close, Egypt and UAE and higher genetic distance with the Turkish population from different continental origin [33] (Table 5). Therefore, North African and Central Asian influences seem to be much higher in Saudi Arabia than in East Asia.

Sample bias-corrected Fst distances were obtained and were represented in multidimensional scaling (MDS) plots (Figure 1). In the plot, Saudis cluster with Middle East populations; Egyptians, UAE and Jewish while Algerian population is found be close to the Middle Eastern population and they are clearly separated from European Turkish; and East Asian populations.

**Fig.1:**
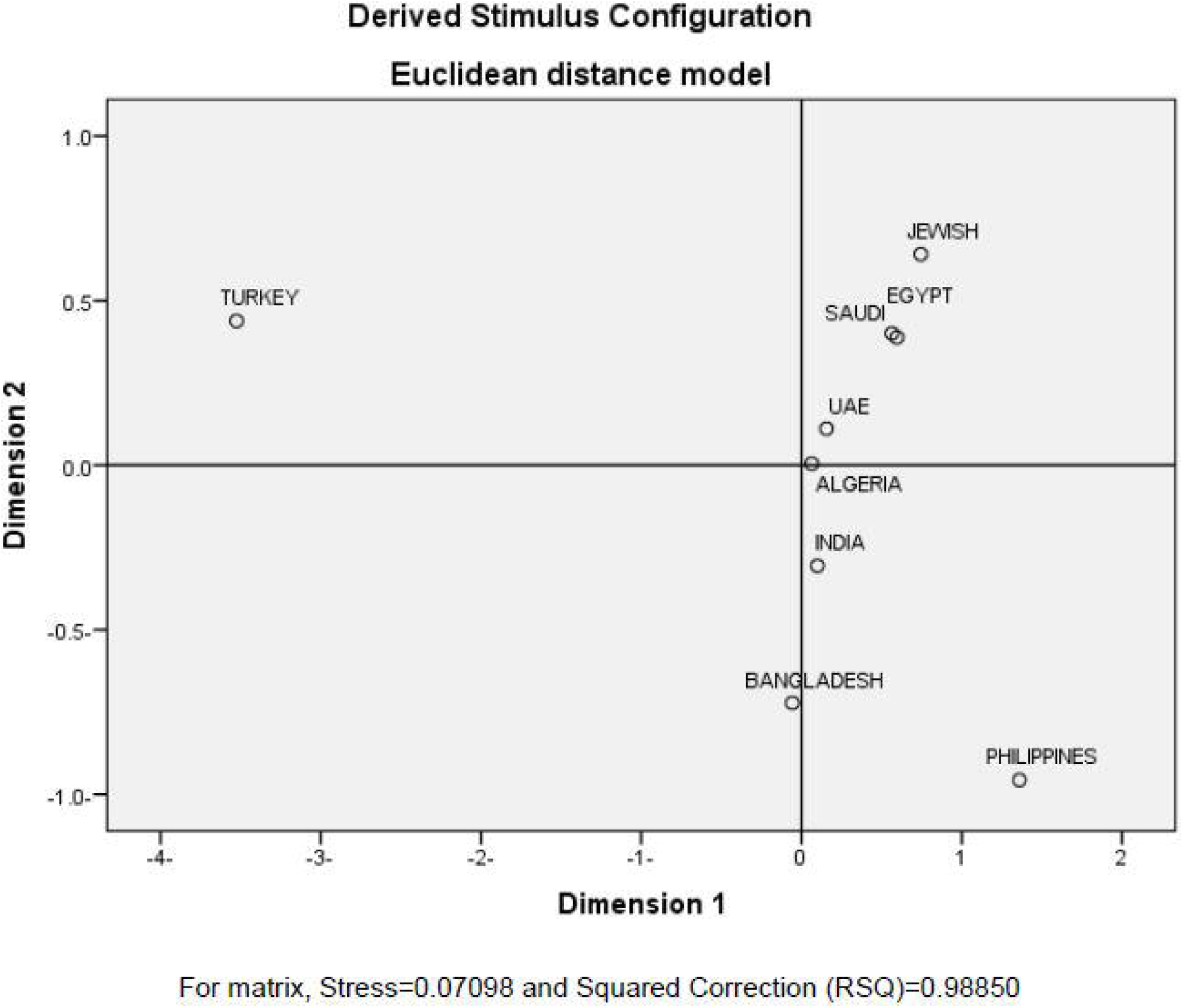
Multidimensional scaling (MDS) plots of the Central Saudi population and other 8 populations [31–38] was built using IBM SPSS Statistics v21.0 software based on the Nei’s genetic distances.

As shown, Saudi, Egyptian, UAE, Jewish and Algerian populations are positioned in the upper right cluster, whereas Indian, Bangladeshi and Filipino populations were clustered together. Turkish population was found in separate clusters apart.

Neighbour-joining (NJ) tree (Figure 2) has also been generated using Nei’s Fst distances of X-STR loci estimated among the 9 populations. This tree graphically represents genetic distance among these 9 populations based on the genotyped 12 X-STR loci. The NJ tree showed that the Saudi population is in close relationship with the Middle Eastern populations (Egyptian, Emirati, and Jewish).

**Fig.2:**
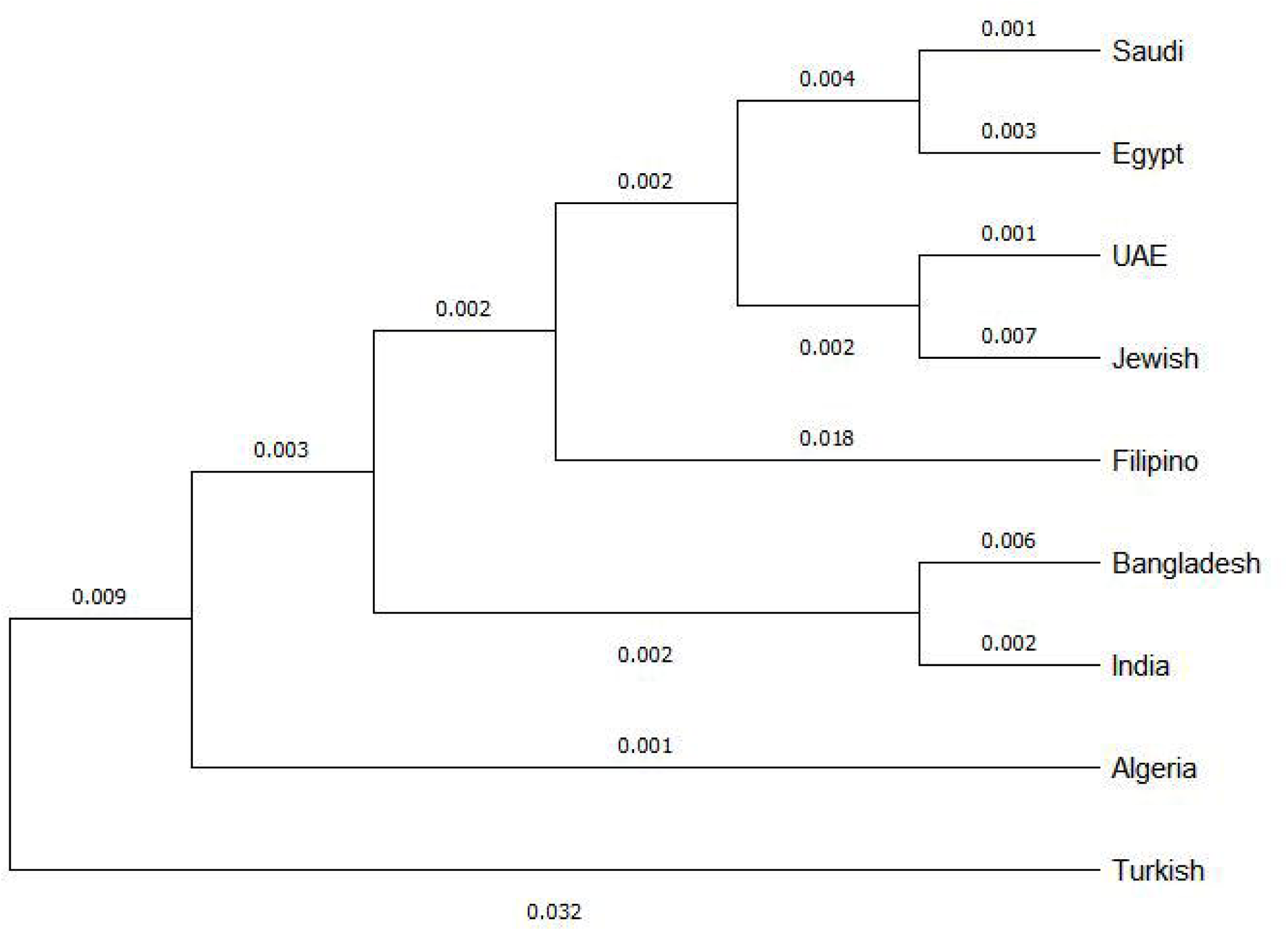
Neighbouring joining tree based on Nei’s Fst Distances for the 12 X-STR loci estimated among the 9 populations [31–38]. The nods are bootstrap values derived from 1000 replications.

The comparison showed a genetic diversity between Saudi population, European and East Asian populations which may be due to geographic and continental origin distinction. Therefore, it is important to develop population data for forensic analysis.

We would like to conclude that the current study was reported to be the first literature exploring the allele frequency and forensic parameters of the 12 X-chromosome STR markers in the Saudi population. The results revealed that X-STRs analysis showed high _c_PD and _c_MECs providing extreme advantages in kinship testing involving female offspring, forensic casework, and human identification. Further, to accomplish the detailed understanding of the genetic structure, intra and inter-population relationships, larger sample sizes must be included from a wider geographic area.

## Acknowledgments

We would like to thank Naif Arab University for Security Sciences for helping to complete this project and allowing us to utilize the forensic Science department laboratories. Also, many thanks to the volunteers for participating in this study.

## CRediT author statement

**Safia Messaoudi**, Conceptualization, Data curation, Formal analysis, Investigation, Methodology, Project administration, Resources, Software, Supervision, Validation, Visualization, Writing – original draft, Writing – review & editing.

**Saranya Ramesh**, Data curation, Project administration, Methodology, Software Writing – review & editing

**Abrar Alsaleh**, Data curation, Methodology, Software

**Mohammed Albajjah**, Methodology, Software

**Noora Alsnan**, Writing – review & editing

**Abdul Rauf Chaudhary**, Writing – review & editing

